# Measuring biodiversity from DNA in the air

**DOI:** 10.1101/2021.07.15.452392

**Authors:** Elizabeth L. Clare, Chloe K. Economou, Frances J. Bennett, Caitlin E. Dyer, Katherine Adams, Benjamin McRobie, Rosie Drinkwater, Joanne E. Littlefair

**Author notes:** **Corresponding author:** Elizabeth L Clare, Department of Biology, York University, Toronto, Ontario, Canada, M3J1P3, **Email:**. **Author Contributions:** ELC designed the experiment and performed field work, CKE performed all laboratory work, FJB performed field work, CED, RD and JEL performed analysis, KA and BM facilitated field work. All authors contributed to manuscript preparation. **Competing Interest Statement:** Authors declare no competing interests. **Classification:** Biological Sciences: Ecology.

## Abstract

Impacts of the biodiversity crisis far exceed our ability to monitor changes in terrestrial ecosystems. Environmental DNA has revolutionized aquatic biomonitoring, permitting remote population and diversity assessments. Here we demonstrate that DNA from terrestrial animals can now be collected from the air under natural conditions, a ground-breaking advance for terrestrial biomonitoring. Using air samples from a zoological park, where species are spatially confined and unique compared to native fauna, we show that DNA in air can be used to identify the captive species and their potential interactions with local taxa. Air samples contained DNA from 25 species of mammal and bird including 17 known (and distinct) terrestrial zoo species. We also identified food items from air sampled in enclosures and detected four taxa native to the local area, including the Eurasian hedgehog, endangered in the UK, and the muntjac deer, a locally established invasive species. Our data provide evidence that airDNA is concentrated around recently inhabited areas (e.g., indoor enclosures) but that there is dispersal away from the source suggesting an ecology to airDNA movement which highlights the potential for airDNA sampling at distance. Our data clearly demonstrate the profound potential of air as a source of DNA for global terrestrial biomonitoring and ecological analysis.

**Significance Statement:** The global decline in biodiversity requires rapid non-invasive biomonitoring tools applicable at a global scale. In this study we collect environmental DNA from mammals and birds from air samples collected in a natural setting. Using only air, we identified 25 species of mammal and bird known to be in the area. Our dataset detected species at risk of local extinction and several confirmed predator-prey interactions. This approach will revolutionize terrestrial biodiversity surveys.

## Main Text

### Introduction

Anthropogenic impacts have caused pervasive biodiversity declines across ecosystems (Díaz et al., 2019; C. N. Johnson et al., 2017; Seibold et al., 2019), particularly from land-use change, habitat loss and degradation (Tilman et al., 2017) leading to the reorganization of global biodiversity patterns and processes (Barlow et al., 2018; Eriksson & Hillebrand, 2019). Rapid and accurate biomonitoring techniques are essential to our attempts to quantify the causes and consequences of global environmental change (Amano et al., 2018; Eriksson & Hillebrand, 2019) and to assist with focused, on-the-ground conservation efforts. Our inability to detect species and measure population dynamics rapidly and accurately is often cited as a fundamental challenge in quantifying our position relative to biodiversity and conservation targets (Amano et al., 2018; C. N. Johnson et al., 2017). Indeed, detecting changes in diversity, abundance, and community composition as well as species range shifts are priorities highlighted by researchers, conservationists, and major international initiatives (Amano et al., 2018). New approaches that provide simpler, large-scale, and automated monitoring techniques are an urgent requirement, needed to address the often-intractable challenge of biodiversity monitoring (Barlow et al., 2018). Decades of development in molecular diagnostics have resulted in established DNA-based approaches for determining species (Blaxter, 2003; Hebert, Cywinska, Ball, & DeWaard, 2003; Tautz et al., 2002) and detecting ecological interactions (Pompanon et al., 2012). The development of DNA reference databases, which permit rapid species identification from unknown environmental samples of (even fragmentary) genetic material, has the potential to transform our ability to monitor global ecosystems. Yet biodiversity monitoring often still relies on capture of live specimens, which is both rate limiting and invasive (Singer, Fahner, Barnes, McCarthy, & Hajibabaei, 2019).

It is well documented that DNA is shed from all organisms and deposited as environmental (e)DNA. This material has been used to analyze contemporary and past ecosystems for nearly two decades (Willerslev et al., 2007, 2014, 2003). An explosion of interest in using aquatic eDNA to assay populations and track invasive species has revolutionized aquatic science, management, and conservation (Ruppert, Kline, & Rahman, 2019). As the field matures, considerable research effort now focuses on the “ecology of eDNA” (Barnes & Turner, 2016) – quantifying and understanding factors influencing eDNA detections beyond inventories alone. Comparative studies have shown that metabarcoding of aquatic eDNA matches or even outperforms conventional methods of community sampling (Bessey et al., 2021; Mena et al., 2021; Ruppert et al., 2019). Additionally, an indication of terrestrial biodiversity can also be obtained from eDNA analysis of water and sediments sampled from aquatic systems (Sales et al., 2020; Ushio et al., 2017), though detections may be biased towards semi-aquatic species. A comparison of tropical mammal detection methods (Mena et al., 2021) including eDNA from lentic and lotic systems, live-trapping, pitfall traps, camera traps, and mistnets found integrated methods provide best estimates of community composition. Although aquatic eDNA alone recovered much of the diversity of mammals (Mena et al., 2021) this would be limited when aquatic systems are not in the vicinity.

A truly terrestrial targeted eDNA system has not yet been developed. On land, eDNA has been measured in permafrost, blood, snow, soil, and honey (Bohmann et al., 2014) and recently by spraying foliage and collecting the runoff to gather eDNA from the surfaces (Valentin et al., 2020). Collecting eDNA from the air, analogous to aquatic eDNA sampling, has remained mostly theoretical (Ruppert et al., 2019) with a few demonstrations recovering DNA from plants (Folloni et al., 2012; M. D. Johnson, Cox, & Barnes, 2019) or fungi (Banchi et al., 2020), although these were based primarily on the analysis of collected dust. We recently demonstrated (Clare et al., 2021) that animal DNA can be extracted directly from air under highly controlled laboratory conditions. The potential for sampling life from air samples could revolutionize terrestrial biodiversity assessments, but to date it has not been tested in the wild. The challenge with validating airDNA methods is establishing an experimental design that permits spatial scales for detection without confounding DNA sources. Zoological parks are ideal for this because they contain captive colonies of mostly non-native species whose identity and spatial location are known with certainty. Indeed, metabarcoding of soils from safari parks, zoological gardens and farms has been used to test the efficacy of these approaches and results have reflected the overall taxonomic richness of terrestrial vertebrates present (Anderson-Carpenter et al., 2011).

Our objective is to use air samples from a zoological park in Huntingdonshire UK to identify zoo species and native wildlife in the first practical application of airDNA sampling under natural conditions. This approach will greatly extend the validation of this technique for global terrestrial biomonitoring and establishes the potential uses of airDNA in ecological systems.

## Materials and Methods

### Methods

#### Sample Collection

This study was conducted at Hamerton Zoo Park, a 25-acre conservation zoo in Huntingdonshire UK established in 1990 and containing approximately 100 species of animal, mostly mammals and birds of conservation concern. It is surrounded by a matrix of agricultural land in rural England. Most species live in enclosures which have free access to outside ranges allowing free air exchange. Air samples were collected using a peristaltic pump (Geotech) and Sterivex-HV filtered (Merck Millipore) with 0.22 µm and 0.45 µm filter sizes. We targeted 15 enclosures which contained zoo species represented in molecular reference collections. For each of these locations, we sampled air for 30 min at 300ml/min filter rate using each filter and we sampled inside an enclosure (e.g. the sleeping chamber) and outside in the open air enclosure (where species move about freely) within 5m of the enclosure opening. We also sampled from general areas of the zoo including the Cat Circle, Owl walkway and near rubbish bins. In addition, we sampled at the Tasmanian Golden Possum Enclosure and Syrian Brown Bear Enclosure, but Golden Possums were not represented in reference databases and the bears were in their hibernation cycle and closed to close sampling (i.e. no indoor samples were taken). We treat these two areas as general areas for sampling. All filters were placed in sterile bags following sample collection and frozen for DNA extraction.

#### DNA extraction

DNA extraction and PCR were carried out within a biological safety cabinet under maximum flow. All extraction procedures followed (Clare et al., 2021). In general, all equipment was sterilized using UV, 10% bleach, 70% ethanol and ultrapure water between each sample. Following existing protocols (Cruaud et al., 2017), the filter was cracked open and the filter removed. DNA was extracted using a Blood and Tissue kit (Qiagen UK) following manufacturer’s protocol but with ATL buffer volumes increased to 450μl to ensure the filters were submerged. We used 50μl proteinase K, 500μl buffer AL and 500μl of 100% ethanol. We used multiple negative controls at the extraction, PCR and sequencing stages. Samples were lysed overnight using a platform shaking at 650rpm at 56°C. The samples were then vortexed and transferred to fresh tubes for extraction. We used QIA shredder spin columns (Qiagen UK) on the remaining filter paper and the flow-through was added to the rest of the sample at which point buffer AL was added. Extraction then followed manufacturer instructions but with centrifugation completed at 11,000 rpm for 3 min following AW2. DNA was eluted in 30 μl of elution buffer pre heated to 70 °C. Elution buffer was cycled through the column three times with 5 min incubation times in each cycle to increase DNA concentration.

#### PCR amplification and sequencing

Each DNA extract was subjected to three PCRs as follows:

*16S PCR* - We amplified a small region of 16S mammal mitochondria using the mam1 and mam2 primers (Taylor, 1996) modified with adaptors for the Illumina MiSeq sequencing platform. The PCR mix included 7.5μl of Qiagen multiplex mix, 1.5μl ddH2O, 5μl of template DNA and 0.5μl of each primer (10μM stocks of each) and amplification used cycling conditions of 95°C for 15min, 40 cycles of 94°C for 30s, 55°C for 90s, 72°C for 90s, and a final 72°C for 10min and a 10°C hold.

*16S nested PCR* - To increase amplification success for low yield sample we performed nested PCRs. For the nested 16S PCRs we first used non-tagged mam1 and mam2 primers. For these reactions, we used 3μl of template DNA (adjusting the amount of water accordingly) and increased the annealing temperature to 59°C. We then used 1μl of each PCR product from the first reaction as a template for a second PCR, again using the same 16S mam1 and mam2 Illumina MiSeq tagged primers. PCR conditions for the second PCR were as previously mentioned.

*COI nested PCR* - We amplified a small portion of the 5’ end of the cytochrome oxidase gene using AquaF2 forward and VR1d reverse primers (Ivanova, Clare, & Borisenko, 2012). We employed a two-step nested PCR strategy. For the first stage PCR, the PCR mix we used comprised 7.5μl Qiagen multiplex mix, 3.5μl ddH2O, 3μl of template DNA and 0.5 μl of each primer (10 μM stocks of each). For the majority of samples, we used 1μl of PCR product from this first reaction as a template for a second PCR using AquaF2 and VR1d Illumina MiSeq tagged primers. For selected samples with significant non-target bands, we gel extracted the target band from the first PCR (Monarch DNA Gel Extraction Kit) and used 1 μl of this purified DNA in the second PCR. Reaction conditions for both first and second PCRs were as follows: 95°C for 15min, followed by 40 cycles of 94°C for 30s, 51°C for 90s, and 72°C for 90s and a final extension at 72°C for 10min, then a hold at 10°C.

#### PCR visualization and sequencing

All products, including positive (cow DNA) and negative controls were visualized using a 1% agarose gel as an initial screening tool and then quantified using Qubit and Tapestation. Amplicons were sequenced on an Illumina MiSeq using unique 5’ forward tags at the Barts and the London Genome Centre following standard protocol using bidirectional 250bp chemistry. The results were demultiplexed by tag for bioinformatics processing.

#### Bioinformatics methods for COI regions

COI read files were uploaded to the mBRAVE platform (http://www.mbrave.net). Paired end samples were assembled with a minimum overlap of 20bp and max substitution of 5bp. Samples were processed to maximize data retention for later steps with the following parameters, Trim Front=38bp, Trim End=26bp, Trim Length=500bp, Min QV filter=0, Min Length=100bp, Max bases with low (<20) QV=75%, Max bases with ultra low QV (<10)=75%. ID threshold=10%, Exclude from OTU at 10% MIN OTU size=1 and OTU threshold=2%.

The reads were compared to the “Hamerton Zoo 1” bespoke reference database consisting of 610 sequences representing 20 species known to reside at the zoo and targeted in our sampling. These data were taken from existing public data in the BOLD database. Sequences not identified by comparison to this bespoke reference collection were then screened in sequential order to system reference libraries:

SYS-CRLCHORDATA (Chordata references) consisting of 40,565 species

SYS-CRLAVES (Aves reference) consisting of 5832 species

SYS-CRLBACTERIA (Bacteria reference) consisting of 2066 species

SYS-CRLFUNGI (Fungi reference) consisting of 565 species

SYS-CRLINSECTA (Insect reference) consisting of 217,994 species

SYS-CRNONINSECTARTH (Non-Insect Arthropoda reference) consisting of 27,832 species

SYS-NONARTHINVERT (Non-Arthropoda Invertebrate reference) consisting of 34,927 species

SYS-CRLPROTISTA (Protista COI reference collection) consisting of 5250 species

#### Bioinformatics methods for 16S regions

We used AdapterRemoval V2 (Schubert, Lindgreen, & Orlando, 2016) to first identify and then remove adapter contamination, using the additional parameters --trimns and --trimqualities, to remove Ns and runs of low quality bases. Read pairs were not collapsed at this step. We processed the remaining reads into amplicon sequence variants (ASVs) using the DADA2 pipeline (Callahan et al., 2016) in R (“R Development Core Team: R: A language and environment for statistical computing,” 2021; RStudio Team, 2020). We filtered the reads using DADA2 with the following parameters: truncate length after 100 bases in both directions (truncLen=c(100,100)), reads with any Ns were removed (maxN=0), reads higher than expected error removed (maxEE=c(2,2)), truncate reads based on low quality scores (truncQ=2) and discard phiX genes (rm.phix=TRUE). Each of the filtered read pairs were dereplicated, the amplicon error rate was estimated, and the core algorithm was used to calculate the true ASVs counts in the data. Finally, read pairs were merged, ASVs in each sample were counted and chimeric sequences were removed.

Final 16S ASVs were blasted against a local subset of the GenBank database (search term: “16S”, downloaded 23rd May 2021, 467,306 records), with >97% identity and output hits limited to 15 sequences. We manually discarded hits with low query coverage (<90%). We then applied BASTA (a last common ancestor algorithm) to the resulting hits, configured to return a majority taxonomy from 90% of the hits (Kahlke & Ralph, 2019). Because the Tyra (*Eira barbara*) was not represented within the 16S reference data we reran this comparison allowing 96% matches to the nearest ancestor in the reference data *Gulo gulo* (not present in the zoo) and assigned ASVs to *Eira barbara* if there was a 96% match to *Gulo gulo*.

#### Data filtering

For both COI and 16S data we excluded *Heterocephalus glaber* or *Fukomys damerensis* identifications as expected contamination from the previous experiment using the same equipment (Clare et al., 2021) and we excluded all human sequences which are expected as a general contamination in all samples and controls. We then examined negative well contamination and recorded identifications in negative samples and the number of reads. We differentiated identifications which would remain if largest negative well ID number was used as a filter and treat each of the three amplifications separately (e.g. a negative well with a 500 reads assigned as a contamination would cause us to flag any ID with 500 or fewer reads assigned, we treat this maximum read count filter separately for COI, 16S and 16S nested PCRs, Supplemental Information).

#### Statistical analysis

Read counts from all three PCR procedures were pooled (Supplemental Information 2) and mean read counts/location for each identified zoo species were calculated. We first examined the effect of the sampling position relative to the animal’s own enclosure (i.e. inside (n=23) or outside (n=79) the animal’s own enclosure) on read counts. In a second model, we examined the relationship between read counts and distance from the animal’s enclosure. The distances between the sampling points to the originating enclosures were calculated as a straight line to the nearest meter using google maps satellite view (i.e. the distance between a sampling point which detected tiger DNA and the tiger enclosure). Distance varied from 0 – 276 metres, but we excluded zero distance datapoints (i.e. datapoints from inside the animal’s own enclosure, n = 23), as this effect had already been examined by the first model. In both cases we used zero-inflated negative binomial mixed effects models using the glmmTMB package (Brooks et al., 2017) in R version 4.0.2 (“R Development Core Team: R: A language and environment for statistical computing,” 2021), with species and filter ID as random effects (filter ID was necessary as we treated read counts from different species from the same filter as different data points). We checked for overdispersion and patterns in the model residuals using the DHARMa package (Hartig, 2021). In both models, we tested the significance of the “sample position” and “distance” terms in explaining the read counts by calculating the likelihood ratio test using the “drop1” function with a chi-squared distribution.

## Results

### Sample collection

We collected 72 air samples from 20 locations around Hamerton Zoo Park. Of these 64 yielded DNA which was identified as belonging to non-human terrestrial vertebrates with multiple sources represented in most samples (Figure 1). All data produced is available on the NCBI short read archive BioProject ID:PRJNA743788.

**Figure 1:**
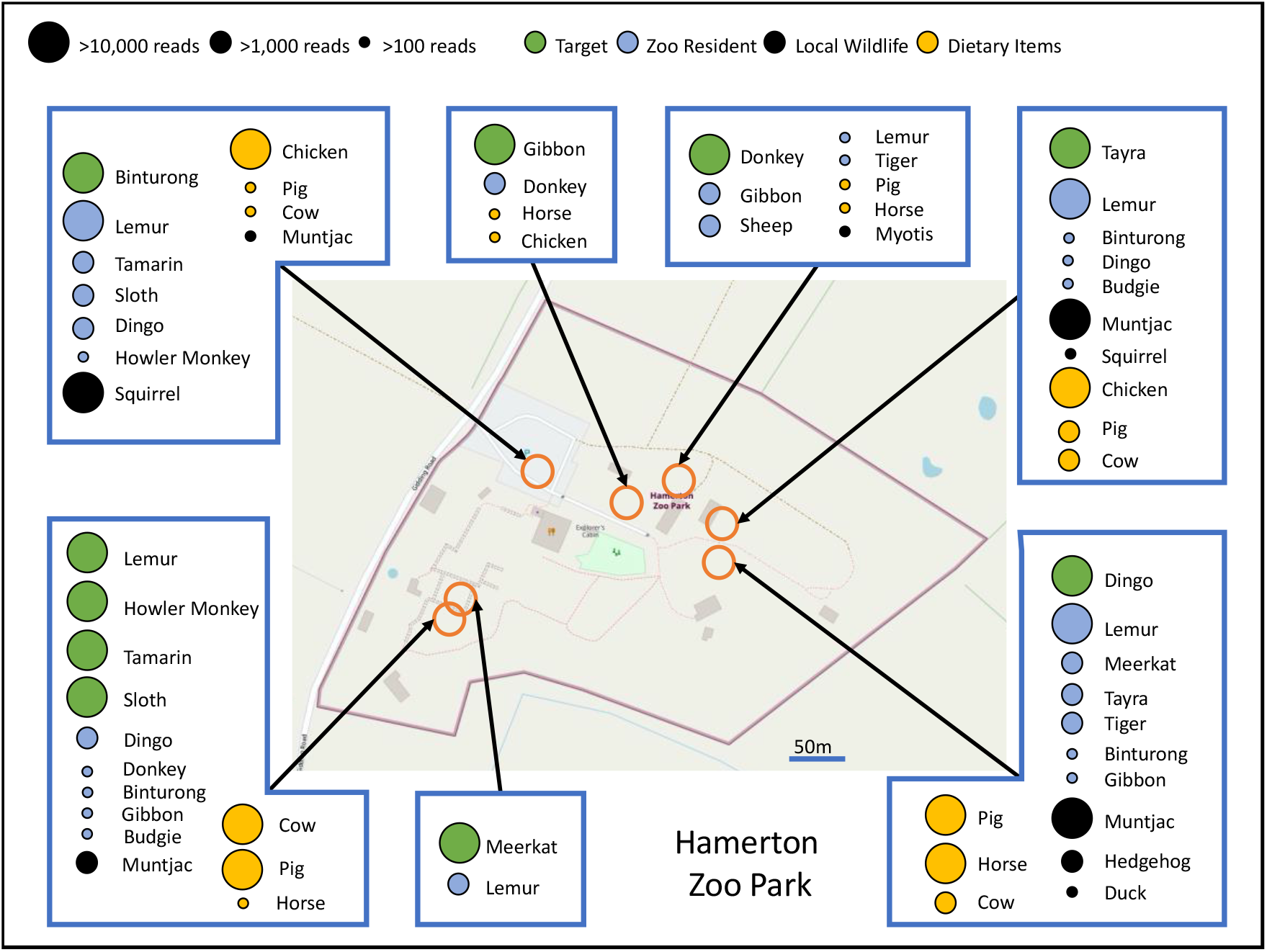
Species identified at seven zoo locations using only DNA collected from air sampling. Identifications are colour coded to indicate the origin of the DNA and circles are scaled to represent approximate read abundance (low, medium and high copy number). Orange rings indicate sampling location. Identifications with <100 copies were excluded from the figure. Full data with read counts for all locations are provided in Extended Table 1, 2 and 3

### 16S Data

We recovered 12,207,070 reads after the removal of adapter sequences; these were used as input into the DADA2 bioinformatic pipeline. After length and quality filtering, paired-end merging and removing chimeras, 11,707,400 reads remained assigned to 335 amplicon sequence variants (ASVs). Taxonomic ID of the ASVs was assigned using BLAST and further refined with BASTA using a last common ancestor (LCA) algorithm and based on 97% sequence similarity to the 16S reference database (see methods for *Eira barbara* identification parameters).

Several ASVs received higher level taxonomic assignments and were resolved as follows. ASVs designated as Artiodacyla were resolved to *Muntiacus reevesi* as the other similar match to a reference was *Cephalophus dorsali* (bay duiker) and is not possible on site. Similarly, ASVs designated a Cervidae were a perfect match to muntjac and a lower match to *Ozotoceros bezoarticus* which was not possible on site. We retain muntjac for these as well. ASVs identified as Herpestidae were perfect or highly similar (>99%) matches to *Suircata suiricata* (which was on site) and lower matches (97%) to other species not present thus we designate these as *S. suricata*. An ASV identified as *Saguinus* was resolved to *Saguinus oedipus* based on matches >99% to that species which was present in the zoo while other potential matches were <98%.

### COI Data

We recovered 6,167,294 reads from samples amplified by COI primers. These data were processed in the mBRAVE pipeline. Filtered data included 1,061,857 reads that were compared to reference databases. From these 361,889 reads were assigned to a non-human mBRAVE BINs (Ratnasingham & Hebert, 2013) at >97% sequence similarity and resolved to species level based on sequence similarity matches >99% in most cases with the exception of *Canis* where species cannot be easily differentiated. We report these as dingo in Table 1, though it is also possible that domestic dog DNA is present on site.

**Table 1:**
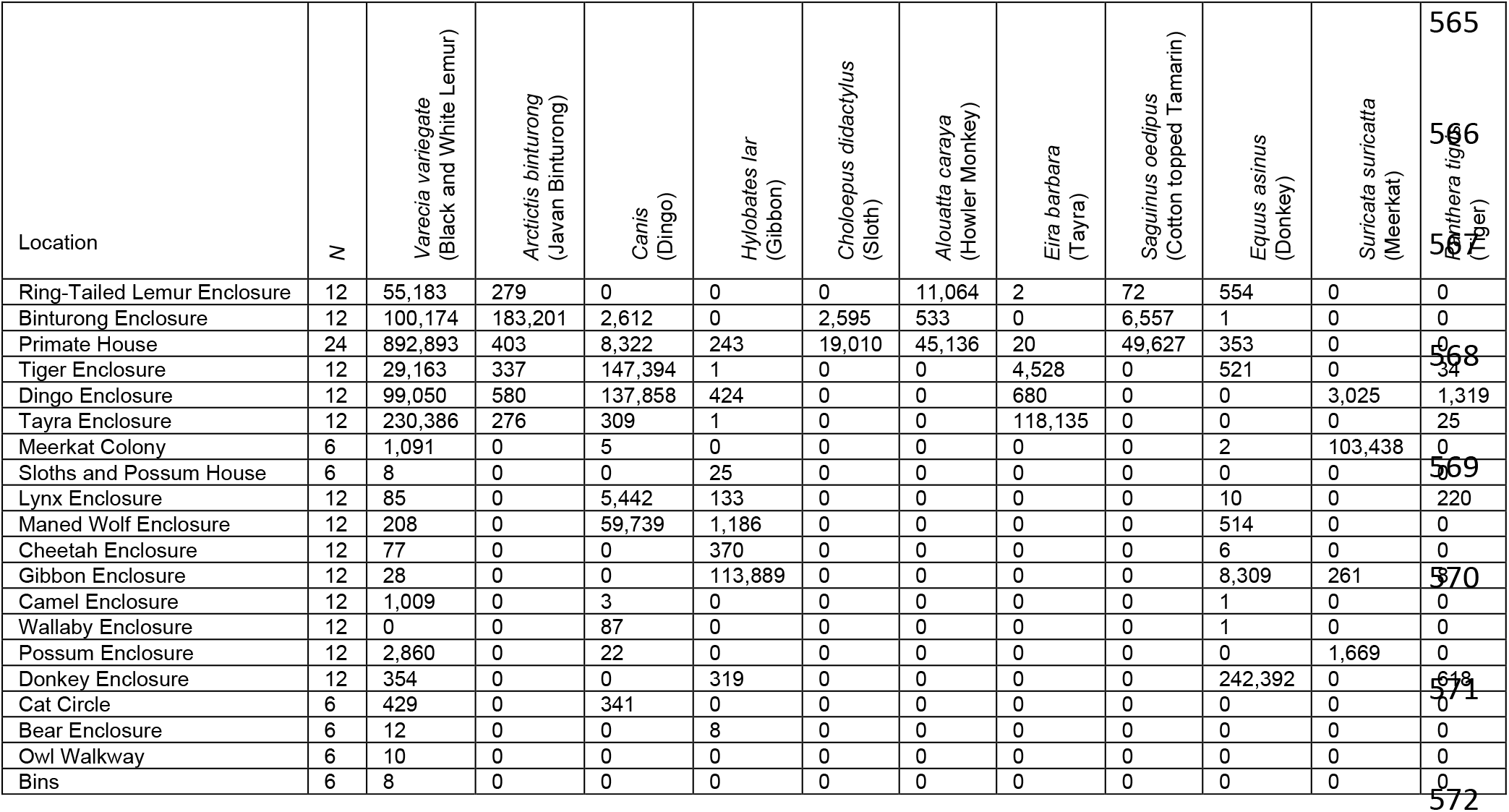
DNA based identification of target zoo species at each location. Cell values represent total read counts from pooled COI, 16s and nested 16s amplifications. Each location was sampled 4 times (inside and outside using 0.25 and 0.45μm filters) with the exception of the Primate House where three inside and one outside space were sampled (eight samples), the Sloth and Possum House which only had an inside space (two samples) and the Meerkat Colony, Cat Circle, Bear Enclosure, Owl Walkway and Bins which were only outside (two samples). N-values represent total number of pooled sequencing runs (samples × 3 PCRs).

### Negative controls for sequence filtering

We used multiple negative controls at DNA extraction, PCR and as empty wells in the sequencing run. Negative well contamination following filtering was very low. However, there was contamination of black and white lemur (1,373 reads) in a negative sample from the 16S PCR negative, donkey (16,836 reads) in a negative of the nested 16S PCR and chicken (513 reads) in a negative of the COI nested PCR. Therefore, in the Supplemental Information for detections, we highlight any read count larger than these to indicate higher support for the taxonomic assignment. Some expected taxa based on sampling location produced read counts lower than these negative thresholds (e.g. *Panthera tigris*) thus we retain all data in Tables 1,2&3 and Supplemental Information to indicate these very likely positives but treat low copy number identifications with caution. All positive control data (cow) was recovered indicating high PCR efficiency and there was very minimal evidence of cow in negative extraction, PCR controls or empty wells used as sequencing controls suggesting that detections in samples represent real dietary detections.

**Table 2:** DNA based identification of non-target zoo species at each location. Cell values represent total read counts from COI, 16s and nested 16s amplifications. Each location was sampled 4 times (inside and outside using 0.25 and 0.45μm filters) with the exception of the Primate House where three inside and one outside space were sampled (eight samples), the Sloth and Possum House which only had an inside space (two samples) and the Meerkat Colony, Cat Circle, Bear Enclosure, Owl Walkway and Bins which were only outside (two samples). N-values represent total number of pooled sequencing runs (samples × 3 PCRs).

| Location | N | Non-Target Zoo Animals |  |  |  |  |
| --- | --- | --- | --- | --- | --- | --- |
|  |  | *Grus carunculata*<br>(Wattled Crane) | *Melopsittacus undulatus*<br>(budgerigar) | *Ovis aries*<br>(Sheep) | *Capra*<br>(Goat) | *Taeniopygia guttata*<br>(Zebra Finch) |
| Ring-Tailed Lemur Enclosure | 12 | 0 | 0 | 0 | 0 | 0 |
| Binturong Enclosure | 12 | 0 | 0 | 0 | 0 | 0 |
| Primate House | 24 | 3 | 443 | 27 | 0 | 0 |
| Tiger Enclosure | 12 | 0 | 0 | 0 | 0 | 0 |
| Dingo Enclosure | 12 | 7 | 0 | 0 | 0 | 0 |
| Tayra Enclosure | 12 | 1 | 442 | 0 | 0 | 0 |
| Meerkat Colony | 6 | 165 | 0 | 0 | 0 | 0 |
| Sloths / Possum House | 6 | 0 | 0 | 30 | 0 | 0 |
| Lynx Enclosure | 12 | 245 | 0 | 0 | 0 | 90 |
| Maned Wolf Enclosure | 12 | 0 | 0 | 0 | 0 | 0 |
| Cheetah Enclosure | 12 | 12,955 | 0 | 0 | 0 | 0 |
| Gibbon Enclosure | 12 | 0 | 0 | 0 | 0 | 0 |
| Camel Enclosure | 12 | 0 | 0 | 231 | 0 | 0 |
| Wallaby Enclosure | 12 | 0 | 0 | 2,791 | 0 | 0 |
| Possum Enclosure | 12 | 0 | 0 | 14 | 0 | 63 |
| Donkey Enclosure | 12 | 1 | 0 | 1,654 | 0 | 0 |
| Cat Circle | 6 | 0 | 0 | 0 | 0 | 0 |
| Bear Enclosure | 6 | 0 | 0 | 231 | 8 | 0 |
| Owl Walkway | 6 | 0 | 0 | 0 | 0 | 0 |
| Bins | 6 | 0 | 0 | 0 | 0 | 0 |

**Table 3:**
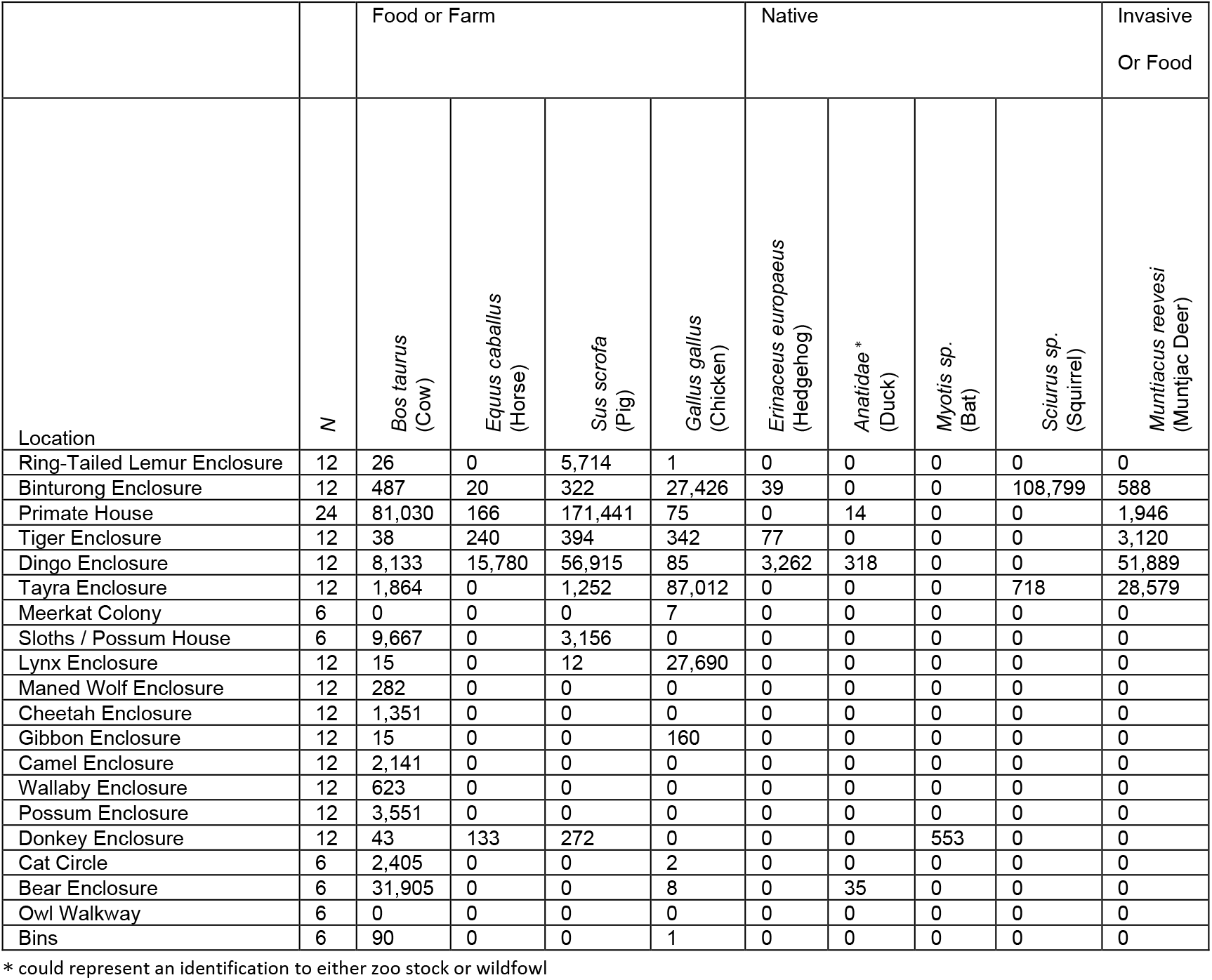
DNA based identification of non-zoo species at each location. Cell values represent total read counts from COI, 16s and nested 16s amplifications. Each location was sampled 4 times (inside and outside using 0.25 and 0.45μm filters) with the exception of the Primate House where three inside and one outside space were sampled (eight samples), the Sloth and Possum House which only had an inside space (two samples) and the Meerkat Colony, Cat Circle, Bear Enclosure, Owl Walkway and Bins which were only outside (two samples). N-values represent total number of pooled sequencing runs (samples × 3 PCRs).

| Location | N | Food or Farm |  |  |  | Native |  |  |  | Invasive<br>Or Food |
| --- | --- | --- | --- | --- | --- | --- | --- | --- | --- | --- |
|  |  | *Bos taurus*<br>(Cow) | *Equus caballus*<br>(Horse) | *Sus scrofa*<br>(Pig) | *Gallus gallus*<br>(Chicken) | *Erinaceus europaeus*<br>(Hedgehog) | *Anatidae\**<br>(Duck) | *Myotis sp.*<br>(Bat) | *Sciurus sp.*<br>(Squirrel) | *Muntiacus reevesi*<br>(Muntjac Deer) |
| Ring-Tailed Lemur Enclosure | 12 | 26 | 0 | 5,714 | 1 | 0 | 0 | 0 | 0 | 0 |
| Binturong Enclosure | 12 | 487 | 20 | 322 | 27,426 | 39 | 0 | 0 | 108,799 | 588 |
| Primate House | 24 | 81,030 | 166 | 171,441 | 75 | 0 | 14 | 0 | 0 | 1,946 |
| Tiger Enclosure | 12 | 38 | 240 | 394 | 342 | 77 | 0 | 0 | 0 | 3,120 |
| Dingo Enclosure | 12 | 8,133 | 15,780 | 56,915 | 85 | 3,262 | 318 | 0 | 0 | 51,889 |
| Tayra Enclosure | 12 | 1,864 | 0 | 1,252 | 87,012 | 0 | 0 | 0 | 718 | 28,579 |
| Meerkat Colony | 6 | 0 | 0 | 0 | 7 | 0 | 0 | 0 | 0 | 0 |
| Sloths / Possum House | 6 | 9,667 | 0 | 3,156 | 0 | 0 | 0 | 0 | 0 | 0 |
| Lynx Enclosure | 12 | 15 | 0 | 12 | 27,690 | 0 | 0 | 0 | 0 | 0 |
| Maned Wolf Enclosure | 12 | 282 | 0 | 0 | 0 | 0 | 0 | 0 | 0 | 0 |
| Cheetah Enclosure | 12 | 1,351 | 0 | 0 | 0 | 0 | 0 | 0 | 0 | 0 |
| Gibbon Enclosure | 12 | 15 | 0 | 0 | 160 | 0 | 0 | 0 | 0 | 0 |
| Camel Enclosure | 12 | 2,141 | 0 | 0 | 0 | 0 | 0 | 0 | 0 | 0 |
| Wallaby Enclosure | 12 | 623 | 0 | 0 | 0 | 0 | 0 | 0 | 0 | 0 |
| Possum Enclosure | 12 | 3,551 | 0 | 0 | 0 | 0 | 0 | 0 | 0 | 0 |
| Donkey Enclosure | 12 | 43 | 133 | 272 | 0 | 0 | 0 | 553 | 0 | 0 |
| Cat Circle | 6 | 2,405 | 0 | 0 | 2 | 0 | 0 | 0 | 0 | 0 |
| Bear Enclosure | 6 | 31,905 | 0 | 0 | 8 | 0 | 35 | 0 | 0 | 0 |
| Owl Walkway | 6 | 0 | 0 | 0 | 0 | 0 | 0 | 0 | 0 | 0 |
| Bins | 6 | 90 | 0 | 0 | 1 | 0 | 0 | 0 | 0 | 0 |

\* could represent an identification to either zoo stock or wildfowl

### Statistical analysis

We compared read counts with distance to most likely source using two models (Figure 2). There was a significant effect of the sample position relative to the animal’s own enclosure on the read counts (Figure 2A, likelihood ratio statistic = 64.1, df = 1, p < 0.001), with read counts inside the animal’s own enclosure being higher (model estimate: 54,899 reads, confidence limits: 19225-156766) than read counts outside the animal’s own enclosure (model estimate: 689 reads, confidence limits: 301-1578). When datapoints from within the animal’s own enclosures were removed (i.e., zero distances), there was no relationship between read count numbers and distance from the enclosure (Figure 2B, likelihood ratio statistic = 3.01, df = 1, p = 0.0828). Neither model was overdispersed.

**Figure 2:**
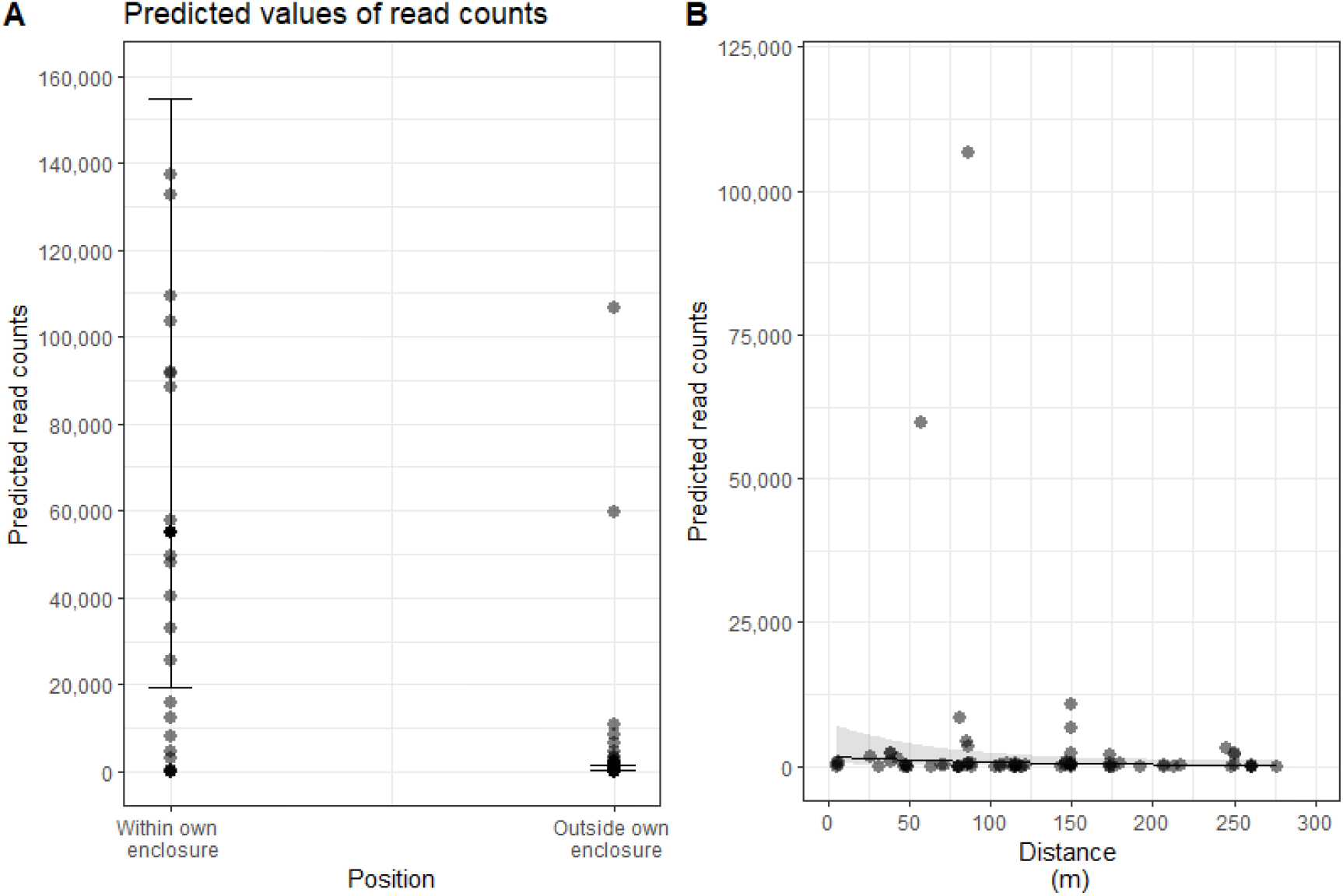
Read count variability with distance from known source. A) Read counts significantly varied according to the sampling position relative to the animal’s own enclosure. Read counts from samples within the animal’s own enclosure were higher than from samples outside the animal’s own enclosure (this also included the enclosures of other animals). B) Read counts were not significantly affected by distance from the animal, once samples from the animal’s own enclosure were excluded. Both plots show predicted read counts from zero-inflated negative binomial models.

## Discussion

Our objective was to collect DNA from air samples and use these to assay for local biodiversity under natural conditions. Using air samples collected at Hamerton Zoo Park we successfully recovered nearly 2.7 million non-human vertebrate DNA sequences. While our laboratory proof of concept (Clare et al., 2021) allowed us to predict that target species could be detected in confined spaces (i.e., inside a sleeping enclosure), detecting airDNA outside enclosures, away from a source, diluted by the air volume in open areas and subjected to wind and local weather represented a far greater challenge. The success of our study shows the potential of conducting biodiversity surveys in the wild, using airDNA, representing an exciting new frontier in biodiversity monitoring.

In addition to the taxa we targeted as part of the known zoo stock, we identified three species of mammal and three species of bird known to be housed at the zoo but in enclosures that we did not have access to (Table 2, Table 3). These additional detections were frequently recovered at the closest sampling point to their actual residence. For example, the indoor exhibit housing budgies, *Melopsitacus undulatus*, and zebra finches, *Taenlopygla guttata*, was closed during the sampling period but we detected their DNA in air samples collected at the adjacent primate house and possum enclosure. While DNA read counts were generally highest within the enclosure where they are expected, we picked up trace read counts in air samples taken more than 250 m from the most likely source (Figure 2). While contamination between samples is theoretically possible, samples were collected and processed on different days and high read counts were retained even after stringent filtering by sequence quality and negative controls. For example, meerkat DNA from an outdoor colony was identified in air sampled at the dingo enclosure 245 m away and at the gibbon enclosure 122 m away. Copy number of recovered sequences was not related to distance from source when enclosures were excluded (Figure 2).

More than a third of the recovered sequences matched cow, horse, pig or chicken. While we cannot preclude DNA drifting in from the surrounding countryside, it is likely these represent food provided to the carnivores. Particularly high concentrations of chicken DNA were detected in the binturong and tayra enclosures while horse, cow and pig were concentrated in samples from the dingo enclosure, so correctly associated with dietary preference (Table 2). Detecting species interactions has been a special focus of environmental DNA approaches (Drinkwater et al., 2019; Pompanon et al., 2012) but this is the first time species interactions have been detected from air. We also observed some unexpected concentrations of these DNA sources perhaps reflecting the movement of people and materials throughout the zoo. For example, an unexpected concentration of pig and cow DNA inside the lemur enclosure could reflect the movement of people between animal houses.

While the primary aim of our study was an inventory of the zoo species, adjacent rural settings are a source for DNA from wildlife *in naturalibus*. We identified DNA associated with squirrels and ducks in several air samples. Several ducks are kept as zoo stock, but we could not identify the genus or species with accuracy, so we classify this as wildfowl but with caution. We may have detected *Myotis* bats, though we also treat this with caution as many bat DNA samples are handled in the processing laboratory facility (Table 2).

Of special interest was the detection of the European hedgehog in three samples. Hedgehogs are commonly observed on site by staff, though they are not as active in the winter thus their detection is particularly interesting. As of 2020, the hedgehog was listed as vulnerable to extinction in the UK (https://www.mammal.org.uk/science-research/red-list/), making it vital to develop additional methods to monitor and protect existing populations. UK species of special interest such as the great crested newt have been the model for the development of aquatic eDNA detection methods (Rees, Baker, Gardner, Maddison, & Gough, 2017) and provide a framework for validating airDNA for similar monitoring. Another commonly cited application of eDNA approaches is the detection of invasive species. We detected muntjac deer, *Muntiacus reevesi*, in five samples. These muntjacs are native to China but became locally invasive after multiple releases in England in the 19th century (Hemami, Watkinson, & Dolman, 2005). They are now well established in the east of England, the location of the zoological park. They are also provided in food for several species on site thus the detection of *M. reevesi* may reflect either food or wildlife (Table 3).

Our study provides compelling evidence that air can be used as a source of DNA for biomonitoring. The detection of multiple taxa in air samples known to reside at the zoo without high false positive detections strongly validates the local source of the DNA. The detection of species of conservation concern and invasive species, as well as DNA from dietary items, likely via the detection of aerosolized fecal material, is compelling and demonstrates the versatility of this genetic approach. High negative rates and low DNA extraction volumes and concentrations suggests a future role in pooling replicate samples, as is done in DNA biomonitoring using leeches (Schnell et al., 2018). This can increase positive hit rates while reducing sequencing costs. The rapid global uptake of aquatic eDNA as a biomonitoring tool highlights the need for new sampling techniques. If airDNA sampling is successfully developed it will have major implications for global terrestrial biomonitoring. The novel opportunities this method provides for tracking faunal composition, non-invasive monitoring of species of special ecological concern, and the detection of species invasion are extremely exciting, and suggest that airDNA could revolutionize the ways in which scientists study and monitor terrestrial biodiversity and could be implemented non-invasively at a global scale.

## Acknowledgments

This study was funded through an EPSRC impact accelerator account provided by Queen Mary University of London (Grant: EP/R511596/1). We appreciate the support of the Hamerton Zoo Park Staff and the Barts and the London Genome Centre for assistance during this project.

## Supplementary Information is available for this paper

Supplementary File 1 contains a detailed table associated with read counts for each PCR reaction.

